# Exploration of inositol 1,4,5-trisphosphate (IP_3_) regulated dynamics of N-terminal domain of IP_3_ receptor reveals early phase molecular events during receptor activation

**DOI:** 10.1101/404020

**Authors:** Aneesh Chandran, Xavier Chee, David L. Prole, Taufiq Rahman

## Abstract

Inositol 1, 4, 5-trisphosphate (IP_3_) binding at the N-terminus (NT) of IP_3_ receptor (IP_3_R) allosterically triggers the opening of a Ca^2+^-conducting pore located ~ 100 Å away from the IP_3_-binding core (IBC). However, the precise mechanism of IP_3_ binding and correlated domain dynamics in the NT that are central to the IP_3_R activation, remains unknown. Our all-atom molecular dynamics (MD) simulations recapitulate the characteristic twist motion of the suppresser domain (SD) and reveal correlated ‘clam closure’ dynamics of IBC with IP_3_-binding, complementing existing suggestions on IP_3_R activation mechanism. Our study further reveals the existence of inter-domain dynamic correlation in the NT and establishes the SD to be critical for the conformational dynamics of IBC. Also, a tripartite interaction involving Glu283-Arg54-Asp444 at the SD – IBC interface seemed critical for IP_3_R activation. Intriguingly, during the sub-microsecond long simulation, we observed Arg269 undergoing an SD-dependent flipping of hydrogen bonding between the first and fifth phosphate groups of IP_3_. This seems to play a major role in determining the IP_3_ binding affinity of IBC in the presence/absence of the SD. Our study thus provides atomistic details of early molecular events occurring within the NT during and following IP_3_ binding that lead to channel gating.

## Introduction

Calcium is a universal and versatile intracellular messenger that regulates a diverse array of biological processes starting from fertilization to cell death^1^. In most cells, generation of calcium signals is initiated by agonist stimulation of cell surface receptors, which activates phospholipase C isoforms, thereby producing inositol 1, 4, 5-trisphosphate (IP_3_) in the plasma membrane. IP_3_ then rapidly diffuses into the cytosol and acts on IP_3_ receptors (IP_3_Rs). The latter represents a major class of intracellular Ca^2+^ channels, present primarily within the membrane of endoplasmic reticulum (ER). IP_3_ binding to the N terminus of IP_3_Rs leads to the opening of the distal pore and the resultant Ca^2+^ flux from the ER lumen elevates cytosolic free Ca^2+^ concentration. IP_3_-evoked Ca^2+^ signals are often spatio-temporally complex and used for regulating various cellular processes. Aberrant activity of these ion channels are known to underlie some pathological conditions^2^.

In mammals, there are three isoforms of IP_3_R (IP_3_R1, IP_3_R2 and IP_3_R3) which share ≥ 60% similarity in amino acid sequence and similar overall domain architecture. A functional IP_3_R is a homo- or hetero-tetramer of subunits, each comprising of ~ 2750 residues. Each subunit consists of a large cytosolic N-terminal region (NT), an intermediary regulatory domain and six transmembrane helices including the pore-forming region towards the C-terminal^3^. Crystal structure of NT of rat IP_3_R1 has been resolved in both *apo* and IP_3_-bound state^4,5^. The NT contains an IP3-binding core (IBC, residues 224-604, rat IP_3_R1) and the ‘suppressor domain’ (SD, residues 1-223, rat IP_3_R1). The IBC can further be envisaged to be made of IBC-β (residues 224-436) and IBC-α (residues 437-604) domains and the cleft formed by these domains accommodate the IP_3_ molecule (Fig. 1). The three NT domains – SD, IBC-β and IBC-α together form a triangular structure where the SD interacts with IBC through two interfaces, β-interface with IBC-β and α-interface with IBC-α. A 310-like turn (α5) extending to the β13 strand of the IBC-β position the IBC relative to the SD, where the SD is located behind the IP_3_-binding site in the IBC. A recent 4.7 Å full length structure of rat IP3R1 solved through cryoelectron microscopy (cryo-EM) ^6^ shows that the IBC is located ~100 Å away from the pore region of the receptor. Quite intriguingly, the cryo-EM structure of rat IP_3_R1 also shows that, unlike the closely related ryanodine receptors (RyRs), the C-terminal tail of the IP_3_R directly interacts with the SD of NT and forms a connecting link between the pore-forming helices and the IP_3_-binding NT region. Very recently, crystallographic analysis also shed light on the long-range communication between IBC and the channel domain. The IP_3_-dependent conformational changes are transmitted to the channel through the three large α-helical domains located between the NT and the transmembrane channel domain^7^. These findings together reaffirm the existence of allosteric modulation of IP_3_-mediated channel gating in IP_3_R.

**Figure 1.**
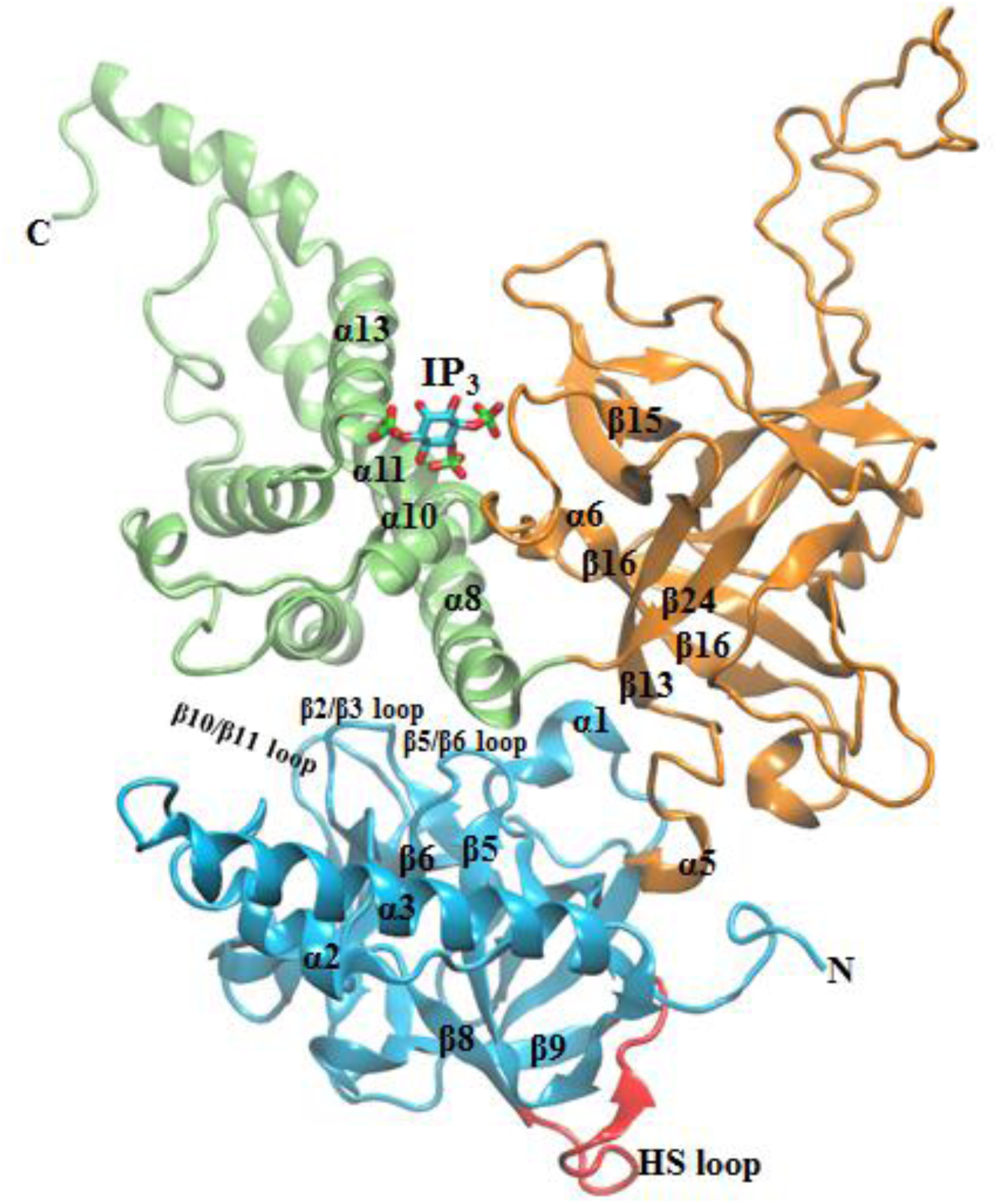
Different sub-domains of IP_3_-bound NT of IP_3_R. The suppressor domain (SD, 1 – 224); IP_3_ binding core-β (IBC-β, 225-436) and IBC-α (437-604) are shown as cyan, orange and green, respectively. The HS loop (residue 165 – 180) of SD that interacts with the IBC-β of a neighboring monomeric subunit in the functional tetrameric form of IP_3_R is colored in red. IP_3_, which binds at the cleft formed between IBC-β and IBC-α is shown in stick representation.

Comparison of the *apo* and IP_3_-bound crystal structures^4,5,7^ provides static snapshots of IP_3_-evoked major domain movements in NT, where the IP_3_ binding brings IBC-β and IBC-α domains close together in a ‘clam closure’ type manner. In agreement with experimental findings^8^, large domain motions in the *apo* IBC as compared to the IP3-bound form, was observed from the molecular dynamics (MD) simulations of the IBC^9^, supporting this ‘clam closure’ during IP_3_ binding. Crystallographic studies further showed IP_3_-induced translational displacement of SD that causes the latter to rotate towards the IBC, measuring an angular displacement of ~ 9 ° between the arm helices (α2/α3 helix, Fig. 1) of the *apo* and IP_3_-bound NT^5^. Such a twist in SD causes translational movement of the conserved HS loop (residues 165-180, rat IP_3_R1) of the SD and the IP_3_ has been proposed to disrupt the inter-subunit interaction between adjacent NTs in the tetrameric IP_3_R that are mediated by IBC-β and HS loop of the SD, allowing the channel to open. Attenuation of IP_3_-evoked Ca^2+^ release in the Tyr167Ala mutant also provided insight into the prime role of the SD in mediating functional coupling between ligand binding and channel gating^10^. However, a dynamic view of the structural events during and immediately following IP_3_ binding at the NT that lead to the opening of a distal (located ~ 100Å away) pore still remains to be elucidated.

As mentioned above, the SD plays a crucial role in traversing IP_3_-binding events at the NT to the channel pore. However, as the name (’*s*uppressor *d*omain’) implies, the SD was found to suppress the IP_3_ binding affinity of IBC^11,12^. Purified protein constructs comprising only the IBC-β and IBC-α binds IP_3_ with higher affinity than the NT or even the whole protein. A cluster of 7 conserved amino acid residues located on one side of the SD were found to be critical for the suppression of IP_3_ binding affinity^13^. IP_3_R isoforms are well known to differ in their IP_3_-binding affinity ^3^ and this has been attributed to the structural difference within their SD^11^. The Gibb’s free energy (ΔG) of IP_3_ binding calculated from the thermodynamics analysis also reflected on the suppressing nature of SD, where ΔG of IP_3_ binding at 296K was −37.1 ± 0.1 kJ/mol and −43.2 ± 0.1 kJ/mol in presence (*i.e.,* NT) and absence (*i.e.,* IBC) of the SD respectively^14^. Suppression of IP_3_-binding affinity of the IBC by the SD is thus well established by now, though the exact molecular basis underlying such phenomenon remains elusive.

In the present study, we have used a computational approach, combining *ab initio* and homology modelling, all-atom MD simulations and principal component analyses to capture the landscape of IP_3_-mediated conformational dynamics in IP_3_R NT and to unearth the molecular details of SD-evoked differential IP_3_ binding affinity of the NT. Analysis of essential dynamics revealed that the binding of IP_3_ triggers a characteristic twist motion of SD, followed by ‘clam closure’ of IBC-β and IBC-α. Inspection of the atomic-level interaction profile led to the identification of a tripartite interaction involving Arg54,Glu283 and Asp444 at the SD-IBC interface, which potentially involved in triggering IP_3_-mediated receptor activation. Very intriguingly, hundreds of nanoseconds long MD simulations of IP_3_-bound IBC in presence and absence of the SD revealed that the ‘Arg269-flipping’ and its positional interaction either with 1^st^ (P1) or 5^th^ (P5) phosphate groups of IP_3_ largely govern the affinity of IBC for IP_3_. To the best of our knowledge, this is the first report that has unveiled the dynamic picture of IP_3_-invoked conformational changes in IP_3_R NT and more importantly, the molecular mechanism underlying the SD-dependent differential IP_3_ binding affinity of the receptor.

## Results and Discussion

To elucidate the conformational dynamics associated with the activation of IP_3_R by the natural agonist IP_3_, we performed all-atom unbiased MD simulations of IP_3_R NT in its *apo* and IP_3_-bound states. Different systems simulated in this study are presented in Table 1 and detailed description of system preparation is provided in the methodology section. Each system was initially simulated for 300 ns of production run following 10 ns of equilibration at 300 K. The results from the simulation studies are discussed below.

**Table 1.**
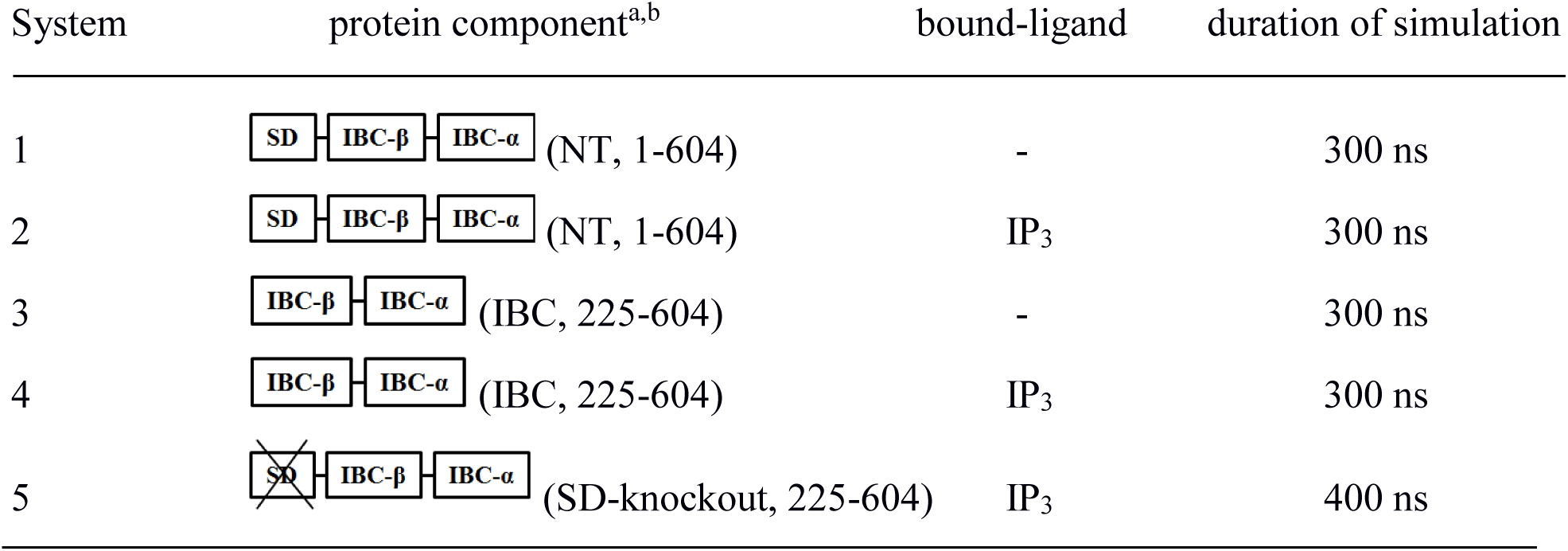
Details of the simulated IP_3_R N-terminal systems ^a^B chain of the crystal structures - PDB ID 3UJ4 and 3UJ0 - were used for the simulation. ^b^System 5 was prepared by removing SD from the final conformation of the IP_3_-bound NT obtained from 300 ns of simulation of system 2

| System | protein component<sup>a,b</sup> | bound-ligand | duration of simulation |
| --- | --- | --- | --- |
| 1 | (NT, 1-604) | - | 300 ns |
| 2 | (NT, 1-604) | IP<sub>3</sub> | 300 ns |
| 3 | (IBC, 225-604) | - | 300 ns |
| 4 | (IBC, 225-604) | IP<sub>3</sub> | 300 ns |
| 5 | (SD-knockout, 225-604) | IP<sub>3</sub> | 400 ns |

### IP_3_ driven twist motion of the suppressor domain

The root mean square deviation (RMSD) of NT with respect to the minimized starting structure as a function of simulation time shows that the IP_3_-bound NT exhibits significant structural deviation, marked by higher RMSDs (Fig. S1a). It is noteworthy that, compared to the *apo* form, IP_3_-bound NT fluctuates considerably throughout the 300 ns of simulation. This observation is further supported by the computed residue-wise root-mean-squared fluctuations (RMSF_s_), indicating increased dynamics of the NT due to the binding of IP_3_ (Fig. S1b). Although RMSD_s_ and RMSF_s_ reflect the conformational changes at NT while IP_3_ binds, it is important to understand the prominent and collective domain motions associated with IP_3_ binding, which can provide insight into the IP_3_-mediated activation of IP_3_R.

We performed principal component analysis (PCA) on simulation trajectories to capture the significant and concerted motions of different NT domains, in presence and absence of IP_3_. PCA identifies the top essential modes (eigenvectors) that represent the major part of collective motions. The top two eigenvectors (principal components, PCs) have captured the significant motions of IP_3_R NT (Fig. S2). To compare the conformational space sampled by *apo* (system 1) and liganded NT (system 2), the two-dimensional projection of the simulation ensembles onto the plane defined by the first two PCs were plotted (Fig. S3a). Although the regions explored by the two systems overlap, notable difference in conformational sampling can be seen between the *apo* and IP_3_-bound states along the PC1 and PC2. A similar trend was also noted from the PCA on the second set of simulations data. This observation is consistent with the results from RMSD and RMSF analyses (Fig. S1a-b), where the protein dynamics was found to be intensified in the presence of the IP_3_.

The principal motions of protein residues can be better visualized and interpreted by representing the eigenvectors as porcupine plots^15^. The porcupine plots showing the motion of SD and IBC of *apo* and IP_3_-bound NT, along the direction of PC1 and PC2, are presented in Fig. 2a-b and 2c-d respectively. In the *apo* state (Fig. 2a-b), loop regions in the NT experienced more flexibility compared to rest of the protein. Particularly, only the arm loop of the SD exhibits significant fluctuations whereas the remaining secondary structures show minimal motions. Similarly, the loop regions of the IBC-β and IBC-α showed comparatively more flexibility than the other part of the domain. Notably, in the *apo* state, the IBC-α exhibits directional movement towards the IBC-β, where the latter experiences negligible mobility (Movie S1).

**Figure 2.**
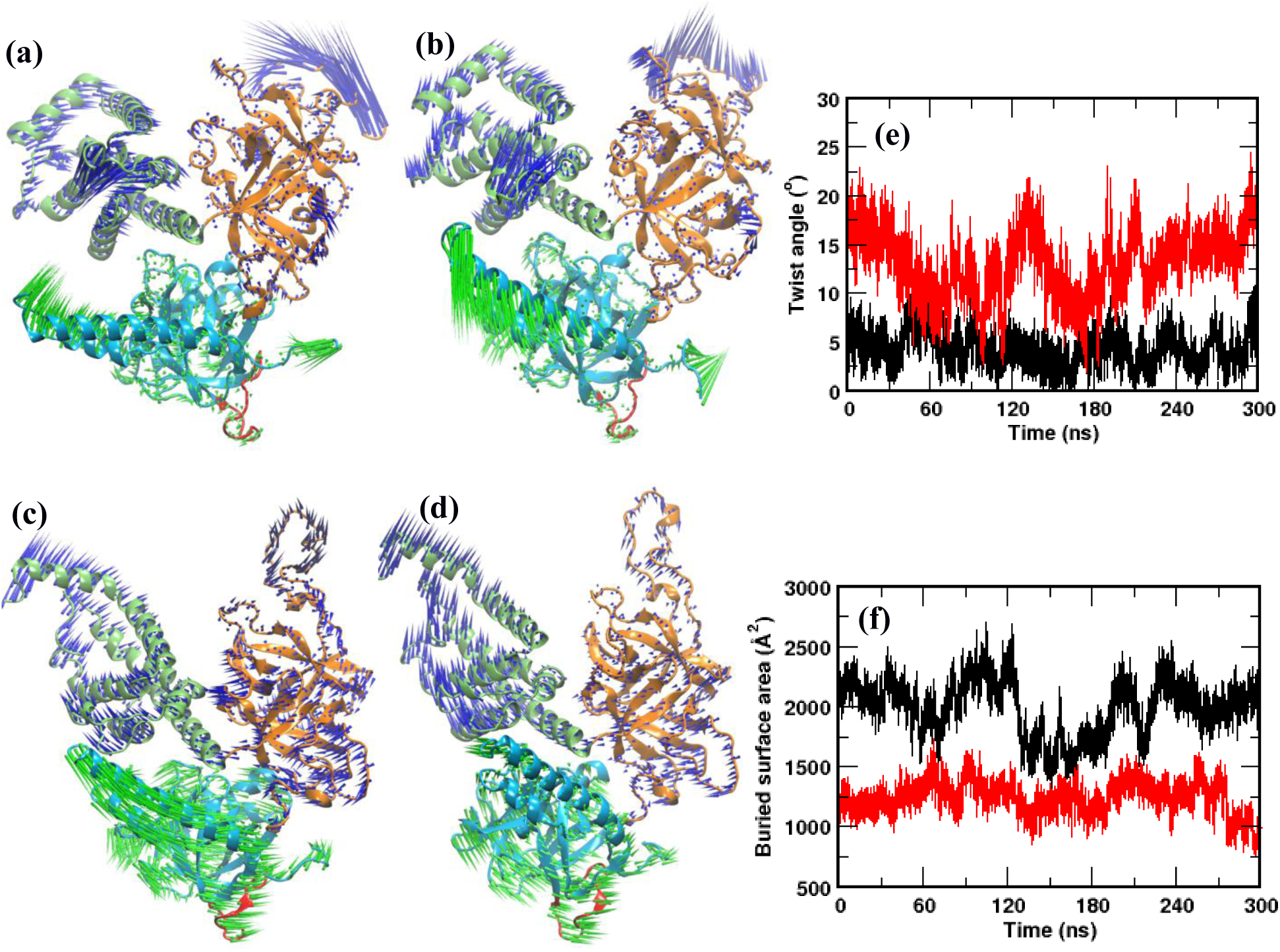
Ligand-induced domain dynamics in IP_**3**_R NT. The porcupine plots representing the principal motions along the direction of PC1 and PC2 respectively are shown for [(a) – (b)] *apo* NT and [(c) – (d)] IP_3_-bound NT. Different domains and the HS loop are colored as shown in Figure 1. (e) The extent of twist motion of the SD measured in terms of twist angle (defined in the methodology) in the *apo* (black) and IP_3_-bound (red) states of IP_3_R NT, as a function of simulation time. (f) A plot of varying SD – IBC buried surface area in *apo* (black) and IP_3_-bound (red) NT, along the simulation time.

Interestingly, IP_3_ binding at the IBC substantially altered the dynamics of different structural units of NT (Fig. 2c-d). The binding of IP_3_ has directly influenced the dynamics of IBC. As in *apo* NT, IBC-α moved towards the IBC-β in the IP_3_-bound NT. Moreover, in contrast to the *apo* NT where IBC-β exhibits minimal displacement, the IBC-β of the IP_3_-bound NT underwent larger dynamics and moved towards the IBC-α. As shown by the vector direction in Fig. 2c-d, IP_3_ induced motions in the NT brings the IBC -β and -α close to each other. This observation is consistent with the findings from earlier studies. The crystallographic experiments showed that IP_3_ binding evokes ‘clam closure’, where the β and α domains of IBC get close to each other ^4,5,8^. Such ligand-induced domain closure is also reported in other ligand-gated ion channels that notably include the ionotropic glutamate receptors^16^. Markedly, as seen from the porcupine plots, the binding of IP_3_ at IBC has allosterically influenced the dynamics of SD. The SD attains a typical twist motion, where the entire SD sweeps over the inner surface of the IBC (Fig. 2 c-d, Movie S2). The SD anchored on the inner surface of the IBC through the IBC-β and the IP_3_ binding causes the SD to twist along the inner surface of IBC-α. The β2/β3, β5/β6, and β10/β11 loops of SD interact with α8 and α9 helices of IBC-α at the α-interface (see Fig. 1). Such a twist motion triggers the SD to move towards the IBC-α, which in turn causes an increased translational displacement of the conserved HS loop of SD. The IP_3_-evoked change in NT dynamics is consistent with the results from RMSD and RMSF analyses, where the increase in RMSD and RMSF in system 2 can be attributed to the IBC clam closure and twist motion of SD (Fig. S1a-b).

The extent of translational displacement of SD due to its twist motion in IP_3_-bound NT is further amplified by calculating the twist angle along the simulation trajectories (described in the methodology). Fig. 2e shows the evolution of twist angle along the simulation time in *apo* and IP_3_-bound systems. It can be noted that the SD twists minimally in the *apo* system with the twist angle fluctuating over a range of 0°-10°, with an average of ~ 5°. In accordance with the observation from essential dynamics (Fig. 2c-d), the angle calculation also captured the increased twist motion of SD caused by the IP_3_ binding at IBC (Fig. 2e). In contrast to the *apo* system, the angle underwent much larger fluctuation (2°-23°, with an average of ~ 13°) during simulation. Such a wide range of twist angle shows the intensive translational displacement experienced by the SD during IP_3_ binding to the IBC. Recent structural studies also support the observed IP_3_ driven twist motion of the SD, where superimposition of *apo* and IP_3_-bound NT revealed ~ 9° angular displacement between the SD arm helices ^5^. However, in contrast to the finding from static crystal structures, MD simulations here provided a dynamic view of IP_3_ mediated binding events at the IP_3_R NT. As depicted in Fig. 2e, the SD twist extensively during IP_3_ binding, where the domain sampled through the conformations with lowest twist angle value of 2°, as seen in *apo* system, as well as explored the conformations exclusive to the IP_3_-bound state, where the SD shows highest degree of twist (~ 23°). The change in buried surface area at SD – IBC interface, illustrated in Fig. 2f, also support the observed twist motion of SD. The computed average buried surface area is 1999 ± 232 Å^2^ in the *apo* system whereas the buried area reduced to 1239 ± 134 Å^2^ in the IP_3_-bound system. Such a reduction in SD – IBC interfacial buried area in IP_3_-bound system implies that the interface has weakened during the IP_3_ binding and many of the residues at the interface are exposed to the solvent. This can be attributed to the enhanced twist motion of SD that affected the SD – IBC interfacial packing. Taking together, the analyses of *apo* and IP_3_-bound systems clearly indicate that the IP_3_ binding at IBC allosterically influences the SD dynamics and the latter experiences a characteristic twist motion.

It is intriguing to note that kinetic analyses of single IP_3_Rs activity recorded using the nuclear patch-clamp technique have shown rapid intra-burst flickers that, unlike the much longer inter-burst intervals, remain largely insensitive to the variation of concentrations of IP_3_ as well as other cytosolic modulators (e.g. free Ca^2+^ or ATP)^17,18^. Such stereotypical, ligand-insensitive flickers were also observed for a mutant rat IP_3_R1 reconstituted in planar lipid bilayers^19^. The flickers were also noticeable, albeit with much lower frequency, in spontaneously active IP_3_Rs recorded from the *Xenopus* oocyte at ultra-low (< 5 nM) level of cytosolic free Ca^2+^ concentration^20^. Based on the PCA and twist angle analysis (Fig. 2), we hypothesise that the basal twist motions observed in the SD in our simulation with the *apo* NT relate to conformations that probably underlie the ligand-insensitive intra-burst flickers of IP_3_Rs, which, however, requires further experimental validation.

### IP_3_ binding breaks the triad of interaction at SD – IBC interface

In the earlier section, we have shown that the SD sweeps over the inner surface of IBC at the SD – IBC interface when IP_3_ binds at the IBC pocket. As observed in earlier structural studies ^4,5,13,21^, our simulations also identified crucial residue contacts at the SD – IBC interface in *apo* and IP_3_-bound NT. Several residues from the SD and IBC involved in interface communication are shown in Fig. S4. A salt bridge between Lys225 and Asp228 positions the SD relative to the IBC and stabilizes the β-interface by helping in arranging the residues at the interface. Notably, a network of hydrophobic and electrostatic interactions *via* Val33, Leu32, Arg54, and Lys127 from SD and Asp444, Phe445, Asp448, Ala449, Val452, and Leu476 from IBC-α are involved in maintaining the α-interface.

Besides identifying residue contacts seen in static crystallographic structures, MD simulations of the NT provided further insight into the allosteric rearrangements in interfacial residue network during IP_3_ binding. The detailed analysis of the *apo* system revealed the formation of a bidentate hydrogen bond (H-bond), involving residues from all the three NT domains – SD, IBC-β and IBC-α. In the *apo* NT, Arg54 of SD forms a bidentate H-bond with Glu283 and Asp444 of IBC-β and IBC-α, respectively (Fig. 3a). It can be noted from the figure that Arg54 is situated at the center of the triangulated structure formed by the three NT domains. The guanidinium side chain of Arg54 H-bonds with the side chain carboxylate group of Glu283 and Asp444, with an average distance of 2.8 Å and 3.2 Å respectively. The -NH group of guanidinium interacts with the -COO group of Glu283 while the guanidinium -NH2 group H-bonds with the -COO group of Asp444. Interestingly, IP_3_ binding at the IBC breaks this triad of interaction at the SD – IBC interface (Fig. 3b). The binding of IP_3_ at the IBC allosterically disrupts the interaction of Arg54 with Asp444 and the former then rearranges its side chain in such a way that the guanidinium -NH2 group H-bonds with the -COO of Glu283. To further evaluate the stability of this bidentate interaction, we calculated the probability distribution of H-bond distance in Arg54 – Glu283 and Arg54 – Asp444 interaction during the entire span of simulation. In *apo* NT (Fig. 3c), Arg54 – Asp444 distance distribution was nearly Gaussian, peaking at the average distance of 2.8 Å, whereas the Arg54 – Glu283 interaction distance was distributed with two peaks, first at 3.2 Å and the second one at 5.1 Å. IP_3_ binding shifts this equilibrium interaction of Arg54 towards Glu283, with the measured Arg54 – Glu283 average distance of 2.8 Å (Fig. 3d). On the other hand, Arg54 was found to be H-bonding with Asp444 (peak at a distance of 2.8 Å) during only about 20% of the simulation time. Moreover, the distribution clearly indicates that, in the *apo* NT, though Arg54 shows comparatively equal affinity towards Glu283 and Asp444, the interaction with Asp444 is found to be stronger than that with Glu283. The energy of Arg54 – Glu283 and Arg54 – Asp444 interaction was calculated as −5.8 ± 2.8 kcal/mol and −15.3 ± 0.6 kcal/mol respectively. Notably, Arg54 – Asp444 interaction energy was decreased to −4.7 ± 0.9 kcal/mol in IP3-bound NT (system 2) while the Arg54 – Glu283 interaction energy was −6.6 ± 1.3 kcal/mol, further supporting the IP_3_-driven residue rearrangements at the SD – IBC interface. The IP_3_ mediated shift in Arg54 – Glu283/Asp444 bidentate interaction can also support the IP_3_-evoked twist motion of SD. In the *apo* state, Arg54 of SD H-bonds with Glu283 and Asp444 and such a bidentate interaction at SD – IBC interface can lock and restrict the conformational flexibility of SD. The IP_3_ binding breaks this triad of interaction by disrupting the Arg54 – Asp444 interaction at the α-interface and stabilises the Arg54 – Glu283 H-bond at the β-interface. Such a residue rearrangement weakens the α-interface allowing the SD to slide over the inner surface of IBC-α and at the same time, anchor the SD immensely on the inner surface of β-interface.

**Figure 3.**
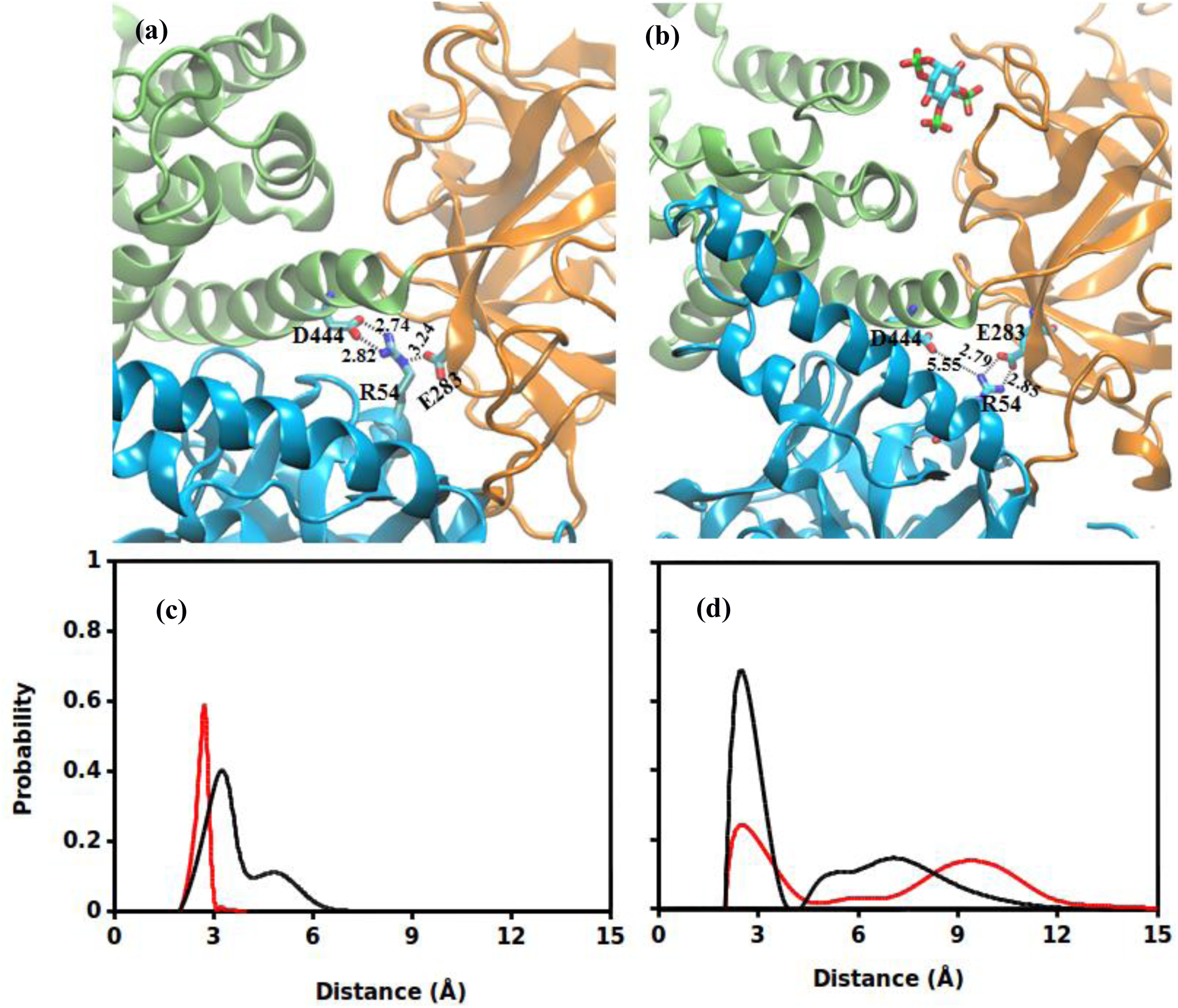
A triad of interactions involving Arg54, Glu238 and Asp444 at the SD – IBC interface found to be crucial for IP3-mediated conformational dynamics. [(a) – (b)] Ensemble averaged structure of IP_3_R NT highlighting the Arg54 interaction with Glu283 and Asp444. (a) Arg54 forms a bidentate interaction with Glu283 and Asp444. (b) IP_3_ binding breaks the Arg54 – Asp444 interaction while maintaining the interaction Arg54 – Glu283. The average distance between the residues is shown. [(c) – (d)] Probability distribution of Arg54 – Glu283 and Arg54 – Asp444 distance in (c) *apo* and (d) IP_3_-bound NT. Distribution is computed over the entire span of simulation for each system. Color scheme: Arg54 – Glu283, black; and Arg54– Asp444, red.

To summarise, the present all-atom MD simulation of IP_3_R NT provides a dynamic view of conformational landscape of NT during and immediately following IP_3_ binding and sheds light on the mechanism of IP_3_-mediated IP_3_R activation leading to the pore opening. In a biologically relevant, functional tetrameric form of IP_3_R, the monomeric units are arranged in such a way that the SD of one monomer interacts with the IBC-β of the neighboring monomer through the HS loop of SD (Fig. S5)^6^. It is proposed that the HS loop – IBC-β interaction holds the tetrameric IP_3_R in a closed state. The IP_3_ binding evokes the IBC clam closure and causes the SD to twist towards IBC, disrupting the HS loop – IBC-β interaction that allosterically opens the ion conducting pore, which is located ~100 Å away from the IP_3_ binding site^6^. In the present study, analyses of essential dynamics of the *apo* and IP_3_-bound NT domains clearly indicate that the IP_3_ binding at IBC allosterically affect the SD dynamics and the latter experiences a characteristic twist motion. The twist motion causes the SD to swing towards IBC-α, which eventually leads to the translational displacement of the conserved HS loop of SD. The residue Tyr167 of the HS loop was found to play a prime role in coupling the IP_3_ binding to gating and mutation of Tyr167Ala had shown to attenuate the IP_3_-evoked Ca^2+^ release^10^. It is noteworthy that, during simulation, Tyr167 experienced a maximum displacement of ~13 Å in the IP_3_-bound system while the displacement was about ~8 Å in the *apo* system (Fig. S6a-b).

### IP_3_ stabilizes IBC domain in the absence of the SD

So far, with the aid of extensive MD simulations, we have explored the IP_3_ driven conformational dynamics in IP_3_R NT. To further investigate the effect of hitherto identified SD twist motion on the IBC dynamics, we simulated two additional systems of *apo* and IP_3_-bound IBC in the absence of the SD (system 3 and 4, Table 1). As the RMSD in Fig. S7 shows, the *apo* IBC (system 3), in the absence of the SD, underwent significant conformational fluctuation from its starting configuration during the course of simulation (average RMSD of 8.4 ± 1.9 Å). However, in the presence of the SD (system 1), IBC gets stabilized after an initial equilibration phase of 40 ns with minimal structural deviation during rest of the simulation (average RMSD of 4.8 ± 1.2 Å). This suggests that, in the *apo* state, IBC experiences conformational constraints in presence of the SD (*i.e.*, as a part of NT, system 1) whereas the IBC becomes increasingly dynamic in the absence of the SD. Interestingly, IP_3_ binding reverses this dynamic behavior of IBC. As shown in Fig. S7, the IP_3_-bound IBC of NT (*i.e.*, in presence of SD, system 2) fluctuates significantly along the 300 ns of simulation, whereas the binding of IP_3_ stabilized the IBC in the absence of the SD (system 4 in Table 1). The RMSD values from the figure clearly indicate that, the liganded IBC is highly dynamic in presence of the SD (average RMSD of 7.1 ± 1.6 Å). On the other hand, in the absence of the SD, RMSD of IP_3_-bound IBC reached a stable plateau after an initial 40 ns of equilibration phase suggesting that the domain is stabilized during rest of the simulation (average RMSD of 5.2 ± 1.4). To further examine the local structural transformations of IBC in the presence and absence of the SD, we compared the residue-wise Cα RMSF of systems 1–4. The calculated RMSF values also corroborate the observation from RMSD analyses. As illustrated in Fig. 4a, in the absence of the SD, the *apo* and IP3-bound IBC exhibit distinct dynamic properties. The IBC residues were comparatively more flexible in the *apo* state (system 3) than the IP_3_-bound state (system 4). However, as seen from RMSD analyses, residue-wise RMSF also shows that the presence of the SD has influenced the dynamic behavior of IBC residues (Fig. 4a, black and red lines). In the presence of the SD (*ie.,* NT), IBC residues experienced increased thermal fluctuations in the IP_3_-bound state as compared to the *apo* state.

**Figure 4.**
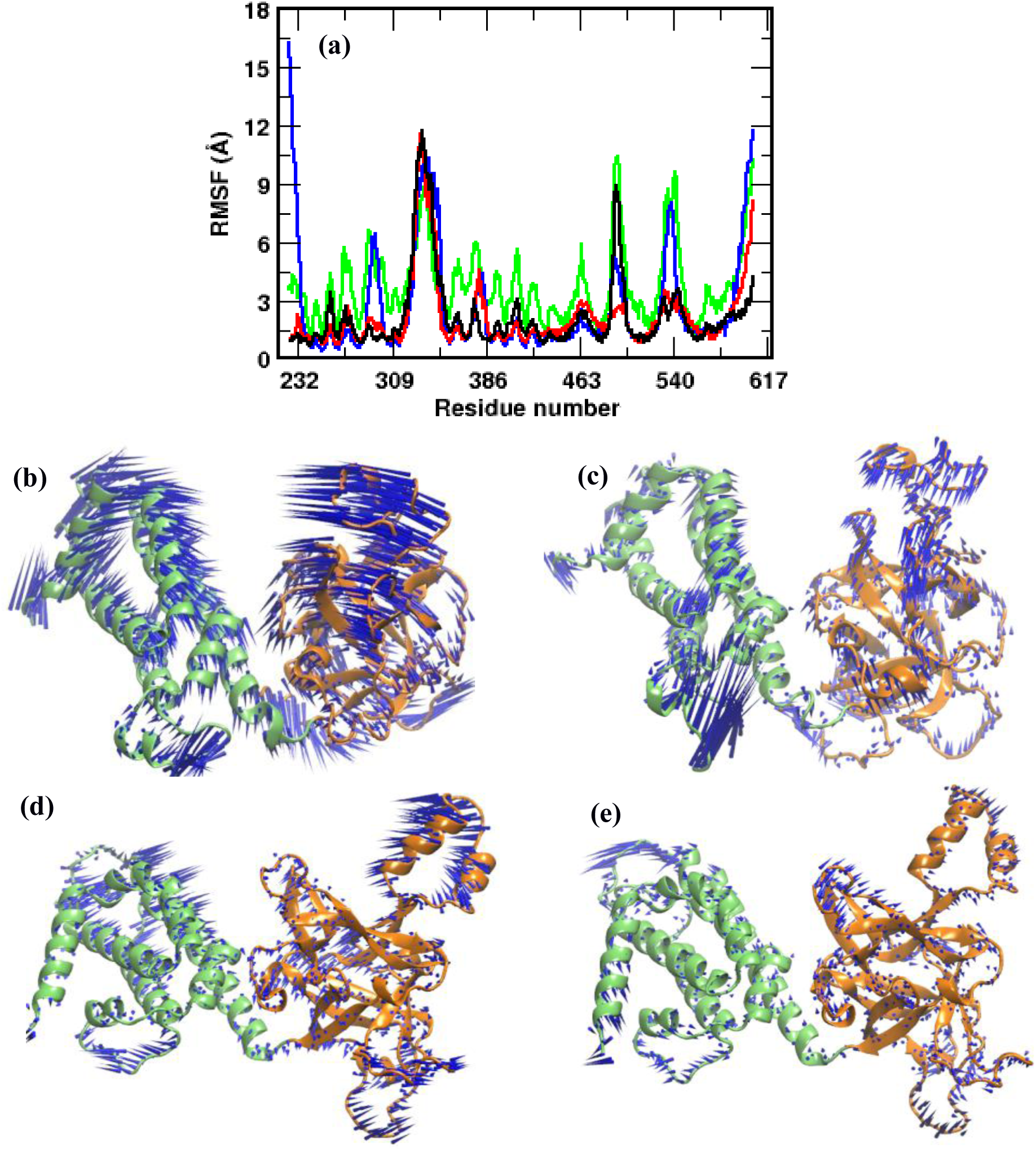
Conformational dynamics of the IBC domains depend on the presence of the SD. (a) Residue-level RMSF of IBC residues in different simulated systems. Color scheme: *apo* NT, black; IP3-bound NT, red; *apo* IBC, green; and IP_3_-bound IBC, blue. The porcupine plots representing the principal motions along the direction of PC1 and PC2 respectively are shown for [(b) – (c)] *apo* IBC and [(d) – (e)] IP3-bound IBC. Different domains are colored as shown in Figure 1.

Furthermore, to understand the domain movements associated with differential dynamics of IBC in the presence and absence of the SD, we resorted to PCA to identify the essential motions in the protein. As shown in Fig. S2, the top two eigenvectors (PC_s_) have captured significant motions of *apo* and IP_3_-bound IBC from each simulation ensemble. The projection of MD ensemble onto the 2D plane defined by the top two PCs plotted in Fig. S3b provides a comparison of conformations sampled by IBC of system 3 and 4. As the figure shows, although conformational overlap is observed, *apo* IBC spans a larger space in contrast to that of the IP_3_-bound IBC. This suggests that the dynamics of IBC is restricted in the liganded state as compared to the *apo* state. This is consistent with the results from the RMSF analysis, indicates that residues with high RMS fluctuations contribute significantly to the dominant motions of the IBC.

The porcupine plots showing the directional motion of *apo* and IP_3_-bound IBC along the direction of PC1 and PC2 in presence and absence of the SD are presented in Fig. 4b-e. Noticeably, in the *apo* state (Fig. 4b-c), all the secondary structure elements of IBC were found to be profoundly dynamic in the absence of the SD (system 3), whereas the IBC exhibits comparatively more restricted motions in presence of the SD (system 1, Fig. 2a-b). In the absence of the SD, *apo* IBC underwent a bending motion where the β- and α-domains of IBC moved towards each other (Movie S3). However, presence of the SD (*i.e*., in apo NT, system 1) has hampered the bending motion of IBC by severely attenuating the dynamics of IBC-β (Fig. 2a-b, Movie S1). Interestingly, IP_3_ binding reverses this characteristic motion observed in the IBC in presence and absence of the SD. In the absence of the SD, binding of IP_3_ has restricted the overall dynamics of IBC (system 4, Fig. 4d-e, Movie S4). The IP_3_ in the binding site prevented the bending motion of IBC domains, which was observed in the *apo* IBC (system 3, Fig. 4b-c). This is expected as the binding site of IP3 located at the IBC-β – IBC-α interface and the bound IP3 imparts local stability by preventing the collapse of the interface that was observed in *apo* IBC. A similar observation was also made from an earlier simulation study with IBC, where the dissociation of IP_3_ conferred larger domain fluctuations to the protein ^9^. However, in contrast to the reduced dynamics of IBC in system 4, IP_3_-bound IBC in presence of the SD (*i.e.*, IBC of IP_3_-bound NT in system 2, Fig. 2c-d) shows increased domain movements, potentially due to the clam closure dynamics and twist motion of SD in the IP_3_-bound NT. The binding of IP_3_ at the IBC allosterically weakens the SD – IBC interface that frees the IBC whereas the stronger SD – IBC interface in the *apo* NT limits the IBC motions. Together, differential dynamics of IBC observed here suggests that a strong inter-domain communication prevails between SD and IBC where both of them influence each other’s dynamic behavior and the binding of ligand have long-range influence on domain dynamics.

### SD suppresses the IP_3_ affinity by shuffling the mode of interaction between the IBC and IP_3_

Though the SD is critical for coupling the IP_3_-induced events to channel-opening, it is well known to suppress the affinity of the IBC or the full IP_3_R protein for IP_3_ binding ^11,12^. The IP_3_ binding affinity of the NT or the whole receptor is reduced by more than one order of magnitude compared to that of IBC alone. Experimentally measured dissociation constants (K_d_) of IP_3_ from different studies showed the higher affinity of IBC for IP3 in the absence of the SD^11,12,14,22^. The binding free energy (ΔG) of IP_3_ at 296K is calculated to be 37.1 ± 0.1 kJ/mol for the NT whereas the IBC, in the absence of the SD, shows higher binding affinity with ΔG of - 43.2 ± 0.1 kJ/mol ^14^. Furthermore, study with different IP_3_R isoforms revealed that IP_3_ binding affinity of the IBC is almost identical in the absence of the SD suggesting the critical role of SD in allosterically modulating the IP_3_ binding to the receptor ^23^. Though previous studies revealed the inhibitory action of the SD, the precise molecular mechanism underlying the reduced binding affinity of IP_3_ in presence of the SD is not yet understood clearly. As discussed in the following sections, the present study with the aid of sub-microsecond MD simulations unveiled the molecular details of ‘suppressing action’ of the SD.

Firstly, to get a qualitative comparison of binding affinity and to complement the experimental observation, we computed the free energy of binding of IP_3_ to NT (*i.e.,* presence of the SD, system 2) and IBC alone (*i.e.,* absence of the SD, system 4) using MM/PBSA approach (see methodology section). The calculated binding ΔG was found to be −70.4 ± 1.2 kcal/mol and −79.5 ± 1.1 kcal/mol for NT and IBC respectively. In agreement with the experimental findings, it is evident from the binding free energy that the IBC, in the absence of the SD, shows comparatively higher affinity for IP_3_ than the whole NT. In order to unravel the molecular mechanism underlying such differential binding affinity, we further analyzed the interaction of individual protein residues with IP_3_ in terms of pairwise interaction energy. The pairwise interaction energy of residues lining the IP_3_-binding pocket is presented in Fig. 5a. Surprisingly, most of the pocket residues interact with IP_3_ with relatively similar strength except Arg269. The interaction energy of Arg269 with IP_3_ was about 15 units more in the absence of the SD as compared to that in the presence of the SD. In presence of SD (system 2), Arg269 – IP_3_ interaction energy was calculated to be −17.3 ± 0.6 kcal/mol. Markedly, in the absence of the SD (system 4), Arg269 of IBC interacts strongly with IP_3_ with energy of −33.8 ± 0.8 kcal/mol. In brief, the energy calculations suggest that the Arg269 – IP_3_ interaction plays a crucial role in determining IP3R’s affinity for IP_3_ in presence and absence of the SD.

**Figure 5.**
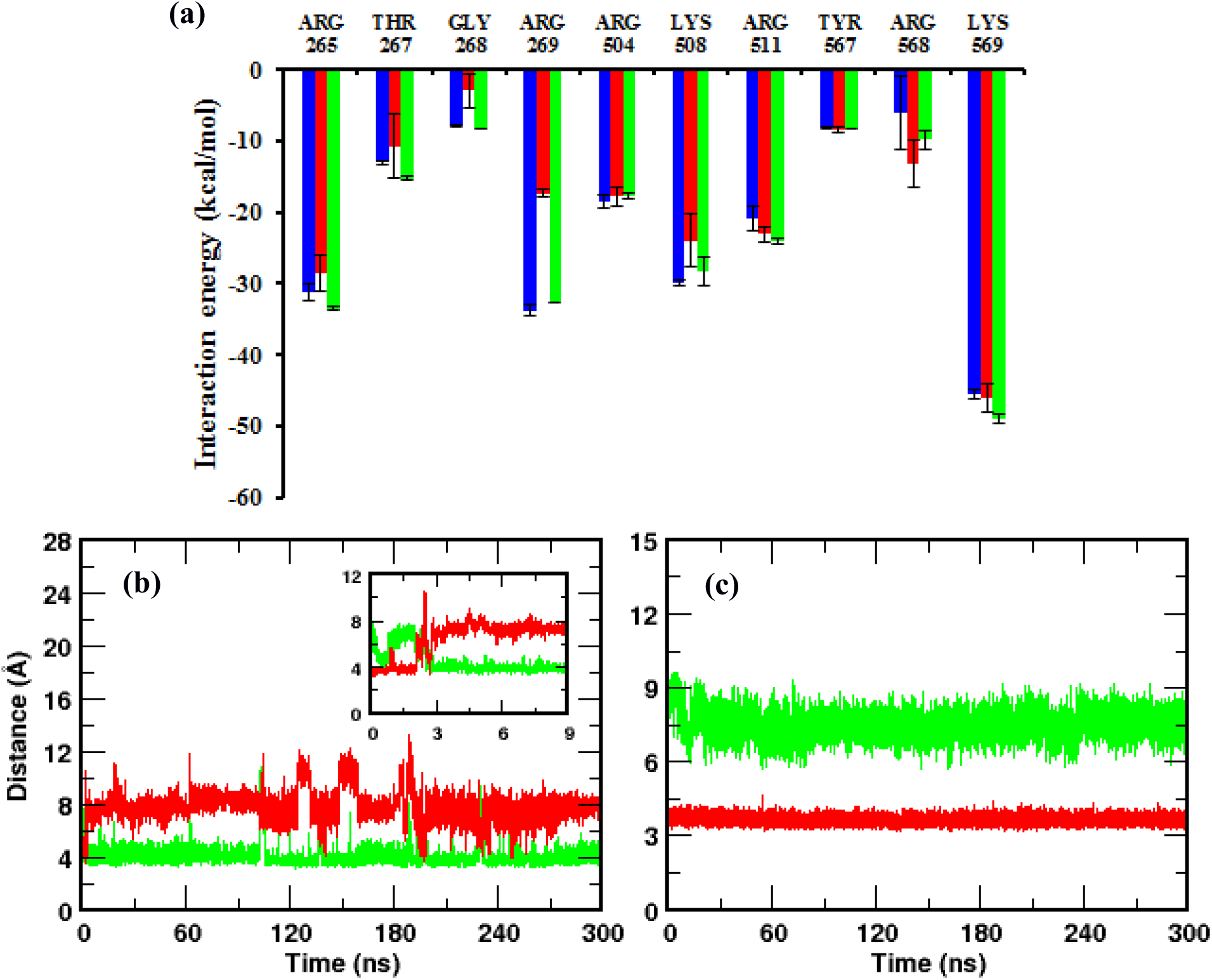
The SD influences the interaction of IP_3_ with the IBC. (a) Residue-level decomposition of interaction energy between IBC pocket residues and IP_3_. Color scheme: IP_3_-bound NT (system 2), red; IP_3_-bound IBC (system 4), blue; and SD-knockout (system 5), green. [(b) - (c)] Time evolution of Arg269 guanidinium – PO_3_ center of mass distance in (b) IP_3_-bound NT and (c) IP_3_-bound IBC. The inset shows the distance in IP_3_-bound NT during the initial 9 ns of simulation. Color scheme: Arg269 – P1, green and Arg269 – P5, red.

To obtain a better understanding of the receptor – IP_3_ interaction, we compared the ensemble-averaged structures of IP_3_-bound NT (system 2) and IBC (system 4) complexes. This is shown in Fig. S8a-b. For clarity, only the ligand and the adjacent protein residues that are involved in direct and consistent interactions are shown. It can be noted from the figure that the sugar moiety and the phosphate groups of the IP_3_ efficiently interact with the residues spanning the IBC -β and -α domains ^24^. As expected, the binding pocket is lined by a number of basic and polar residues to accommodate a highly negatively charged molecule like IP_3_. Three phosphate groups - P1, P4, and P5 - of the IP_3_ molecule are involved in an extensive network of H-bonds with five arginine residues - Arg265, Arg269, Arg504, Arg511 and Arg568. The Lys508 that lies at the interior of binding pocket also forms H-bond with the P5 of IP_3_. During the entire span of simulation, Thr267 and backbone amino group of Gly268 was found to interact with P4 phosphate group. The C6’-OH of the IP_3_ forms H-bond with either Arg504 or Tyr567 in both system 2 and system 4, whereas other hydroxyl groups fail to make any direct interaction with protein residues. Such a less significant role of IP_3_ hydroxyl groups in its binding with IP_3_R is reported from the biophysical studies with IP_3_ derivatives^25^.

As discussed above, most of the pocket residues show similar pattern of interaction with IP_3_ in the presence and absence of the SD. However, very interestingly, the SD seemed to influence the mode of interaction of Agr269 with IP_3_. In the presence of the SD (system 2), the guanidinium group of Arg269 was found to interact effectively with the P1 (phosphate group attached to the C1’ of the inositol sugar ring) of the bound IP_3_ molecule (Fig. S8a). On the other hand, in the absence of the SD (system 4), Arg269 of the IBC makes multiple interactions with the bound IP_3_, where the guanidinium group of the residue H-bonds with P5 (phosphate group attached to the C5’ of the inositol sugar ring) and C6’-OH of the IP_3_ molecule (Fig. S8b). In both systems, the guanidinium -NH/-NH_2_ groups were involved in H-bond with the 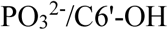 groups of the IP_3_ molecule. As evident from the figure, Arg269, which interacts with the P1 of the IP_3_ in the presence of the SD (*ie.,* in NT), is repositioned in the absence of the SD and forms H-bond with the P5 and C6’-OH groups of IP_3_ that lies deep inside the pocket. We further examined the stability of Arg269 – IP_3_ interaction by measuring the 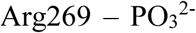 - distance along the 300 ns simulation trajectories (Fig. 5b-c). In the presence of the SD, the guanidinium group of Arg269 interacts with the P1 having a H-bonding distance of ~ 3.8 Å throughout the simulation whereas the P5 was found to be located ~ 8 Å away from the Arg269 (Fig. 5b). Strikingly, Arg269 flipped its mode of interaction in the absence of the SD (system 4). As shown in Fig. 5c, rather than interacting with the P1 (distance of ~ 7 Å) as seen in the presence of the SD, the guanidinium group of the residue H-bonds with P5 in the absence of the SD, maintains a distance of ~ 3.8 Å. A similar analysis of time evolution of Arg269 – C6’-OH distance during simulation is depicted in Fig. S9. The guanidinium -NH_2_ – C6’-OH average distance was measured to be 5.8 ± 1.0 Å in the IP3-bound NT (system 2) as compared to the H-bond distance of 3.9 ± 0.5 Å in the IP3-bound IBC (system 4).

A closer look at the initial configurations from the simulation trajectories has also provided further insight into the SD-invoked Arg269 flipping. In the crystal structures, Arg269 was found to interact with the P5 of the IP_3_ in both IP_3_-bound IBC (PDB ID: 1N4K) as well as NT (PDB ID: 3UJ0) with a distance of about 2.7 Å. It is to be noted that the region spanning residues 268-270 that includes Arg269 is not resolved in any of the apo-NT crystal structures (PDB IDs: 3UJ4, 3T8S apo, 5X9Z, 5XA0) as well as an IP_3_-bound NT structure (PDB ID: 3T8S). It is, however, apparently resolved in the IP_3_-bound NT structures of the Cys-less rat IP_3_R1 (PDB ID: 3UJ0) ^5^ as well as the mouse IP_3_R1 (PDB ID: 5GUG, 5XA1)^7^. But, for all these structures, the crystal structure of mouse IP_3_R1 IBC (PDB: 1N4K) that is clearly devoid of the SD, effectively served as the reference for the initial molecular replacement^5,7^. We suspect that this region containing Arg269 is very flexible and this is supported by our simulation as well as close inspection of the available crystal structures of the NT and full length IP^4,5,7^. Intriguingly, within the initial 3 ns of the simulation in our study, Arg269 within the IP_3_ bound NT breaks its interaction with P5 and flips towards P1, measuring a reduction in Arg269 – P1 distance from 8 Å to ~ 3.8 Å (Fig. 5b inset) and retained the distance of ~ 3.8 Å during rest of the simulation. It can be postulated that, in the starting NT configuration, the SD influenced the overall IBC architecture where the loop bearing the Arg269 was unable to occupy in its initial conformation, forcing the Arg269 to reposition itself to interact with the P1 of the IP_3_.

To strengthen our finding further that the SD modulates Arg269 flipping and interaction with IP_3_, we created a new simulation system (system 5) by removing the SD from the final conformation of IP_3_-bound NT obtained after the 300 ns simulation of system 2. Hence, the newly generated system consists of IP_3_-bound IBC of the previously simulated NT where the Arg269 interact with the P1 of the IP_3_. Here, our aim was to examine whether the IBC can sense the absence of the SD and reorient the Arg269 in such a way that it interacts with P5 as seen in IP_3_-bound IBC (*i.e.,* in the absence of the SD, system 4). As in the above section, we analysed the Arg269 – IP_3_ interaction in system 5 by monitoring the guanidinium 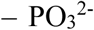 distance along the simulation time. In Fig. 6, we combined the calculated distance from system 2 and 5. To our surprise, IBC in system 5 senses the absence of the SD during the course of simulation that eventually leads to the necessary structural rearrangements in the IBC with flipping of the Arg269 towards P5. As shown in Fig. 6, after ~ 280 ns of simulating the SD-removed NT (*i.e.,* IBC), Arg269 – P5 distance decreased from about 10 Å to the H-bonding distance of ~ 3.8 Å and, at the same time, Arg269 – P1 distance increased to about 8 Å. This suggests that Arg269 breaks its interaction with P1 and flips towards P5 after ~ 280 ns of simulating the SD-removed IP_3_-bound IBC. Remarkably, the flipping of Arg269 was also reflected in the IP_3_ binding affinity. As expected, the IP_3_ binding affinity of IBC for IP_3_ was increased in the absence of the SD, with a rise in ΔG of binding from −71.9 ± 2.1 kcal/mol, in the IP_3_-bound NT, to −80.9 ± 0.9 kcal/mol, after the removal of the SD. The flipping has also altered the Arg269 – IP_3_ interaction energy (Fig. 5a). By shifting the site of interaction from P1, in presence of the SD, to P5, after the removal of the SD, Arg269 has strengthened its interaction with IP_3_ with an increase in energy from −17.3 ± 0.6 kcal/mol to −32.6 ± 0.1 kcal/mol. It is noteworthy that the calculated energies are in good agreement with that calculated from IP_3_-bound IBC in system 4 (Fig. 5a, blue bar). A movie showing the flipping mechanism of Arg269, in the absence of the SD, in system 5 is presented in Movie S5. Together, these analyses suggest that the SD allosterically modulates the IBC interaction with IP3 and Arg269 plays a crucial role in determining the binding affinity of IBC for IP_3_ in presence and absence of the SD. Higher IP_3_ binding affinity in the absence of the SD can be ascribed to the multiple interactions of Arg269 with IP_3_, involving Arg269 – P5 and Arg269 – C6’-OH, in contrast to the Arg269 – P1 interaction in the presence of the SD (Fig.5b-c, Fig. S8 and Fig. S9).

**Figure 6.**
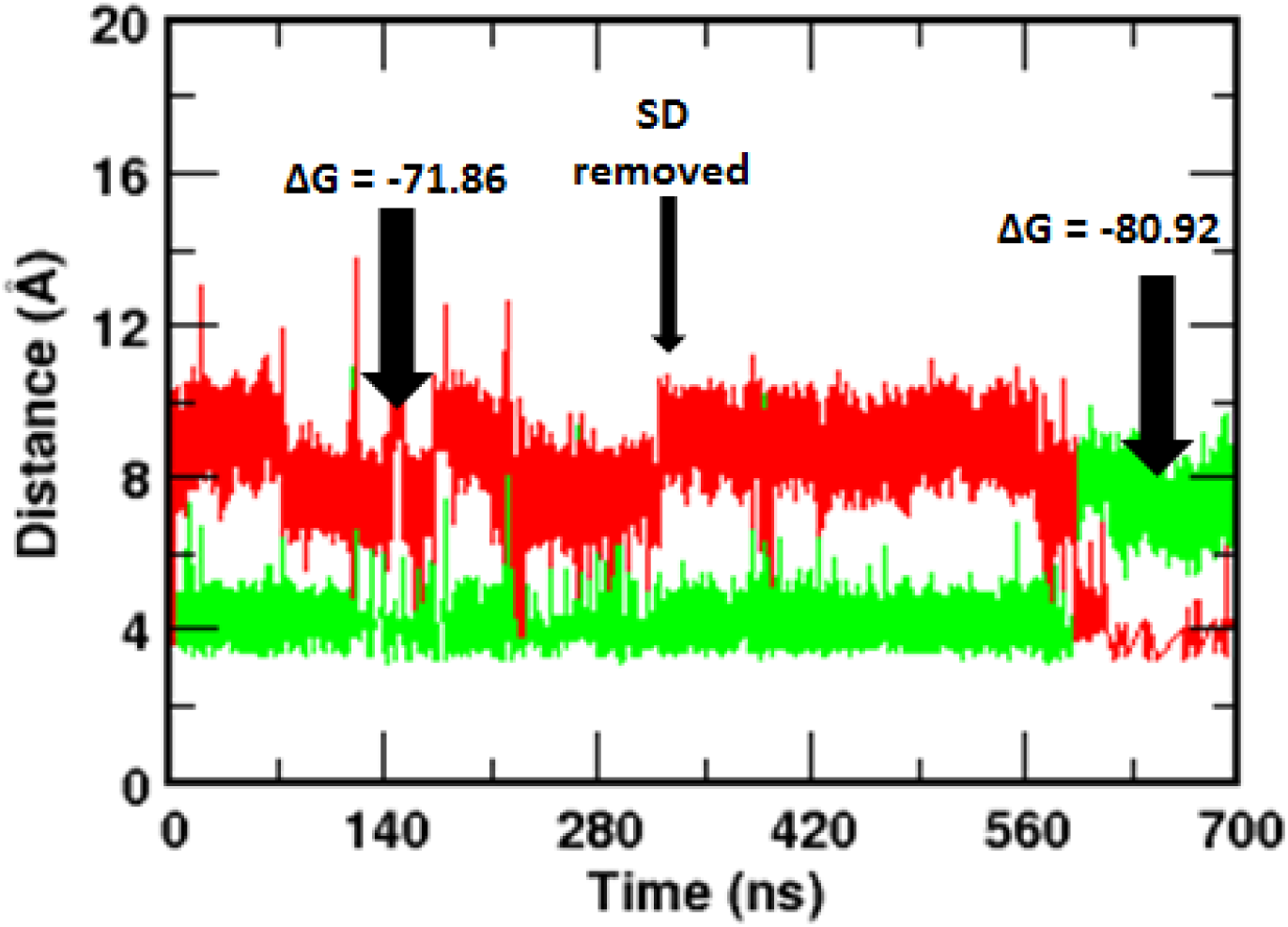
Removal of the SD from the IP_3_-bound NT flips the Arg269 interaction with the IP_3_. Time evolution of Arg269 guanidinium – PO_3_ center of mass distance combined from system 2 and system 5 (see Table 1 for system definition). The SD was removed from IP_3_-bound NT (system 2) after the initial 300 ns of simulation and thus generated IP3-bound IBC (system 5), which was simulated for another 400 ns. Corresponding free energy of IP_3_ binding before and after the removal of SD are shown. Color scheme: Arg269 – P1, green and Arg269 – P5, red.

## Conclusions

As with most other ligand-gated ion channels, a detailed understanding of the activation gating of IP_3_Rs remains a holy grail in the field. Although, the currently available structural and biochemical information has significantly advanced our understanding of the structure-to-function relationship in IP_3_R, details of the dynamics of IP_3_-mediated domain movements within IP_3_R NT remained largely unexplored. In the present study, extensive MD simulations of *apo* and IP_3_-bound NT and analysis of essential dynamics have provided a detailed picture of correlated domain movements during and immediately following IP_3_ binding. Compared to the *apo* form, SD of IP_3_-bound NT showed characteristic twist motion, where the SD move towards the IBC-α. During the binding process, IP_3_ has brought together the β and α domains of IBC, leading to the ‘clam closure’. Our results demonstrated a dynamic view of unique structural plasticity acquired by the IP_3_R NT as a consequence of IP_3_ binding and support the observations inferred from crystallographic studies^5^. The H-bond and energetic analyses revealed the IP_3_-induced molecular rearrangements at the SD – IBC interface, where Glu283-Arg54-Asp444 triad of interaction breaks during the IP_3_ binding. The Arg54-Asp444 interaction breaks while the Arg54 maintains its interaction with Glu283. MD simulations further show the existence of inter-domain dynamic correlation in the IP3R NT and reveal that the SD is critical to the conformational dynamics of IBC. Very interestingly, the study unraveled the SD-dependent flipping of interaction of Arg269 between P1 and P5 phosphate groups of IP_3_, which seems to play a major role in governing the IP_3_ binding affinity of IBC in the presence/absence of the SD.

To conclude, the present study has brought out the importance of alternate conformations of IP_3_R NT needed for its function and the comprehensive understanding of the IP_3_-binding mechanism of IP_3_R derived here could help to understand the physiological activation and pharmacological regulation of IP_3_Rs.

## Materials and Methods

A series of unbiased all-atom MD simulations of IP_3_R NT (residues 1-604) in its *apo* and IP_3_-bound states (Table 1) were performed. The crystal structure of the IP_3_R NT was used to model the full length NT with residues 1-604. Thus, the atomic coordinates of the *apo* (PDB ID: 3UJ4) and IP_3_-bound (PDB ID: 3UJ0) IP3R NT obtained from the protein data bank were used in this study. The missing residues; 1-6, 76-84, 204, 269-270, 294-298, 373-381, 407, 426-427, 486-500, 525-550 and 578-604 were incorporated with the help of MODELLER9v16 ^26^ using crystal structures of IBC (PDB ID:1N4K) and SD (PDB ID: 1XZZ) as templates. Additionally, the crystal configurations were also missing a sequence of residues 319-351. Since we noted that a very low sequence similarity with available structures in databases, the 3D model of this sequence was constructed using the *ab initio* protein structure predictor, QUARK ^27^. Quark builds the models based on structures of small fragments of 1-20 residues, which was later assembled by replica exchange Monte Carlo simulations using an atomic-level knowledge-based force field. The modeled structure of the amino acid sequence 319-351 was merged with the crystal conformation of the *apo* (system 1) and liganded NT (system 2) and optimized by the conjugate gradient energy minimization method in Modeler program ^26^. Further, the *apo* (system 3) and IP_3_-bound (system 4) IBC-only structures of IP_3_R NT were generated from the full length NT by deleting the N-terminal SD of residues 1-224.

In each of the above protein structures, hydrogen atoms were added using H++ server ^28^, maintaining the ionization state of the residues at the pH of 7.4. A set of partial atomic charges for IP_3_ was obtained via quantum calculations. A B3LYP geometry optimization procedure was performed using Gaussian 09 with the 6-311+G* basis set ^29^. The atom-centered RESP charges were calculated via fits to the electrostatic potentials obtained from the calculated wave functions^30^. The interaction parameters for the IP_3_ were adopted from a previous work, where the forcefield parameters for phosphorylated inositols were developed based on OPLS-AA/AMBER framework^31^. Subsequently, the structures were energy minimized for 1000 steps using the steepest descent algorithm and another 1000 steps by the conjugate gradient method, using AMBER 14.0 simulation package ^32^. After relaxing the added atoms in gas phase, each structure was solvated in cubic periodic box of explicit water with water molecules extending 14 Å outside the protein complex on all sides. The 3-site TIP3P water model was chosen to define the water molecules. In system 1 and 2, the simulation box contained ~ 45,000 water molecules, while ~ 22000 water molecules were present in the systems 3 and 4. Charge neutrality was maintained by adding one Cl-ion in system 1 and five, three and four Na^+^ ions in system 2 - 4 respectively. The solvated systems were subjected to extensive energy minimization and thermalization with raising the temperature gradually to 300 K in canonical ensembles, maintaining harmonic restraints on crystallized heavy atoms of protein and ligand. Afterwards, solvent density was adjusted under isobaric and isothermal conditions at 1 atm and 300 K and the harmonic restraints were slowly reduced to zero. To enable the volume variation, simulations were performed in an NPT ensemble using the Berendsen thermostat and barostat. The systems were equilibrated for 10 ns, with a 2 fs simulation time step. The long-range Coulombic interactions were treated using Particle Mesh Ewald sum with a cut off of 12 Å applied to Lennard-Jones interactions and all bonds involving hydrogen atoms were constrained using the SHAKE algorithm. Each system was later subjected to 300 ns of production run, saving the trajectories at an interval of 2 ps for further analysis. All simulations were performed using the AMBER 14.0 molecular dynamics simulation package with the AMBER ff14SB force field ^32^. Additionally, a new simulation system (system 5, Table 1) was generated from the final conformation of IP_3_-bound NT obtained from 300 ns of simulation of system 2. This ‘SD-knockout’ system was prepared by removing the N-terminal SD of 224 residues from the IP_3_-bound NT. Like other systems, simulation of system 5 was performed for 400 ns following a similar protocol as stated above. Together, a total of 1.6 μs of atomistic simulations were carried out in this work.

Principal component analysis (PCA) is employed to explore the dominant motions of *apo* and IP_3_-bound IP_3_R NT region. PCA tool has been used in the past to locate the essential motions in complex systems with high dimensionality, like in proteins, and the detailed methodology can be found elsewhere^33^. PCA was performed in this study using pyPcazip program package^34^. The eigenvectors from PCA were visualized as porcupine plots using VMD^35^, where the direction of the porcupine corresponds to the direction of the motion and the length of the porcupine denotes the magnitude of the motion. The twist of the SD was calculated by measuring the angle defined by the two structures, the crystal structure and a snapshot from the simulation. A plane was defined on each structure by three points that is located on the center of mass of each domain (SD, IBC-β and IBC-α). The ‘twist angle’ defined as the dihedral angle between the two planes was calculated. PyMOL (The PyMOL Molecular Graphics System, Version 1.7.4, Schrodinger, LLC) and VMD^35^ was used to generate all structural figures.

*Free Energy Calculations –* Binding free energies of IP_3_ were calculated using Molecular Mechanics Poisson Boltzmann Surface Area (MMPBSA) method implemented in Amber tools package^32^. The block averaged ΔG values were calculated from five independent windows of 5ns generated by taking the last 25 ns trajectory and each window has 100 snapshots from the trajectory. In general, the free energy of ligand binding to protein in solvent is estimated as follows,

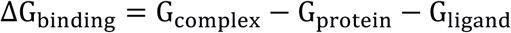

where G_*complex*_, G_*protein*_, and G_*ligand*_ are the free energies of the protein-ligand complex, unbound protein, and free ligand, respectively. The ΔG_*binding*_ can also be calculated from the changes in the molecular mechanical gas-phase energy (ΔE_*MM*_), entropic contribution, and solvation free energy:

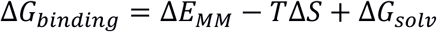

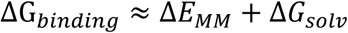

where ΔE_MM_ is the vacuum potential energy that considers the bonded and non-bonded interactions between the protein and ligand. The solvation free energy, ΔG_*solv*_, is estimated by solving the linearised Poisson Boltzman equation for each of the three states (ΔG_*polar*_) and adding an empirical term for hydrophobic contributions to it (ΔG_*nonpolar*_). The hydrophobic contribution is calculated from the solvent accessible surface area. As in practice, the entropic contribution (*TΔS*) was omitted since the calculations involve binding of similar ligand to the same protein. Therefore, the computed values will be termed as the relative binding free energies.

### Data availability statement

Most of the data generated or analyzed during this study are included in this article (and its Supplementary information files).

## Acknowledgments

This work was supported by a Royal Society research grant and Royal Society University Research Fellowship (to T.R.). A.C. was a Royal Society Newton Fellow. D.L.P. was supported by the Biotechnology and Biological Sciences Research Council UK. X.C. was a recipient of the Cambridge Trust-IDB Malaysia scholarship. AC thank Prof. Saraswathi Vishveshwara (Molecular Biophysics Unit, Indian Institute of Science Bangalore, India) for critically reading the manuscript.

## Author Contributions

A.C. and T.R. conceived the project. A.C. performed the research and analysed data with input from X.C. and D.L.P. A.C. and T.R. wrote the paper.

## Additional Information

### Supplementary information

9 supplementary figures and 5 supplementary movies accompanies this paper.

